# SALMA: Scalable ALignment using MAFFT-Add

**DOI:** 10.1101/2022.05.23.493139

**Authors:** Chengze Shen, Baqiao Liu, Kelly P. Williams, Tandy Warnow

## Abstract

Multiple sequence alignment is essential for many biological downstream analyses, but accurate alignment of large datasets, especially those exhibiting high rates of evolution or sequence length heterogeneity, is still unsolved. We present SALMA, a new multiple sequence alignment that provides high accuracy and scalability, even for datasets exhibiting high rates of evolution and great sequence length heterogeneity that arises from evolutionary processes. Like some prior methods (e.g., UPP, WITCH, and MAFFT-sparsecore), SALMA operates in two distinct stages: the first stage computes a “backbone alignment” for a subset of the sequences, and the second stage adds the remaining sequences into the backbone alignment. The main novelty in SALMA is how it adds the remaining (“query”) sequences into the backbone alignment. For this step, which we refer to as SALMA-add, we use divide-and-conquer to scale MAFFT-linsi--add to enable it to add sequences into large backbone alignments. We show that SALMA-add has an advantage over other sequence-adding techniques for many realistic conditions and can scale to very large datasets with high accuracy (hundreds of thousands of sequences). We also show that SALMA is one of the most accurate compared to standard alignment methods. Our open source software for SALMA is available at https://github.com/c5shen/SALMA.

## 1 Introduction

Multiple sequence alignment (MSA) is a crucial precursor to many downstream biological analyses, such as phylogeny estimation [16], RNA structure prediction [20], protein structure prediction [9], etc. Obtaining an accurate MSA can be challenging, especially when the dataset is large (i.e., more than 1000 sequences), has a high evolution rate, or has sequence length heterogeneity. Some of these issues have been reasonably well addressed through the newer alignment methods, but sequence length heterogeneity, whether resulting from large insertions or deletions (i.e., “indels”) in the evolution of the sequences or through the inclusion of reads or incompletely assembled gene sequences, still presents significant challenges for accuracy [17, 22, 26]. Various techniques have been developed for this problem, with UPP [17], WITCH [21] and MAFFT-sparsecore [27] perhaps the most relevant. Each of these methods, albeit with slight differences in design, operates in two stages: the first stage extracts and aligns a subset of the sequences, and the second stage adds the remaining sequences into the constructed subset alignment. Of the three methods, WITCH has been shown to improve on UPP for accuracy but is more computationally intensive [21], and no comparison has yet been made between MAFFT-sparsecore and these other methods.

A related computational problem of increasing importance is adding one or more sequences into an existing alignment. This problem is relevant to the case, for example, of an evolutionary biologist interested in constructing large gene family trees and then updating them rather than computing the alignment from scratch each time a new sequence is made available. There are specific commands within MAFFT that enable sequences to be added into a user-provided alignment, including the --addfragments and --add commands [11]. Moreover, the UPP and WITCH methods also explicitly enable sequence addition into existing alignments, by allowing the user to provide a backbone alignment and thus skip the first stage of their two-stage process (this capability does not seem to be available in MAFFT-sparsecore). Thus, these two MSA problems are related, with solutions for the second problem (i.e., adding sequences into an alignment) relevant to the first (i.e., computing MSAs in the presence of sequence length heterogeneity).

We present SALMA, a new approach to aligning sequences with sequence length heterogeneity. Like UPP and WITCH, SALMA operates in two stages where the first stage extracts and aligns a subset of the sequences, and the second stage adds the remaining sequences into that initial subset alignment (which we will refer to as the “backbone alignment”). The innovation in SALMA is in its second stage, where it uses a new divide-and-conquer technique, referred to as SALMA-add, to add the remaining sequences into the backbone alignment. Moreover, this approach enables SALMA to align datasets with hundreds of thousands of sequences with high accuracy. We show, through an experimental study involving large biological and simulated datasets (both nucleotides and proteins) that SALMA scales to large datasets. We study alignment of two domains from serine recombinases, which perform numerous biological roles: replicon resolution, DNA inversion, and chromosomal integration of genomic islands. These proteins are often encoded on mobile DNAs, so expected to have a high rate of evolution. It is also one of the most accurate alignment methods under many biologically realistic conditions.

The rest of the study is structured as follows. We present the background material, including a discussion of existing methods, in Section 2, and we present SALMA in Section 3. We present the design of the experimental study in Section 4, and the results of the study are provided in Section 5. We discuss trends in Section 6 and conclude with directions for future work in Section 7.

## 2 Background

### 2.1 UPP-add

We refer to the second stage of UPP [17] as UPP-add. Given a backbone alignment, UPP-add builds an ensemble of Hidden Markov Models (i.e., an eHMM) to represent the backbone alignment, which it then uses to add query sequences into the backbone alignment. To build the eHMM, UPP-add computes a backbone tree on the backbone alignment using the maximum likelihood heuristic FastTree2 [19]. Then, UPP-add recursively decomposes the backbone tree into smaller subtrees by deleting “centroid edges” until the last decomposition results in subtrees with at most *A* leaves (*A* = 10 in default UPP). Each subtree thus defines a subset of the backbone sequences and hence also a sub-alignment (induced by the backbone alignment on the subset of sequences). An HMM is created using *hmmbuild* from the HMMER suite [6] for each sub-alignment, which produces the eHMM. Each query sequence is searched against all HMMs in the eHMM using *hmmsearch*, mapped to the HMM with the highest bit-score, and then added into the sub-alignment using *hmmalign*. Because the sub-alignments are all compatible with the backbone alignment, this also defines a way of adding the query sequence into the backbone alignment. The set of all query-backbone alignments are merged using transitivity to form the final alignment.

### 2.2 WITCH-add

WITCH-add (i.e., the second stage of WITCH) improves upon UPP-add by considering more than one HMM to align each query sequence [21]. WITCH-add computes a backbone tree and creates an eHMM in the same way as UPP-add and performs an all-against-all search between all query sequences and all HMMs. Then, it uses the bit-score from the all-against-all search to calculate a weight between each query-HMM pair, and *k* top-weighted HMMs are used to align each query sequence (*k* = 10 in default WITCH). Then, it uses Graph Clustering Merger (GCM), a merging technique from MAGUS [25], to obtain a merged alignment for each query sequence from their corresponding query-HMM alignments. Finally, all merged query alignments are added to the backbone alignment transitively in the same way as UPP-add. As shown in [21], WITCH-add recovers more true homologies than UPP-add, at the cost of a minor increase in false homologies.

### 2.3 MAFFT-add

MAFFT-add [11] in its default setting uses a standard progressive alignment procedure with two iterations to add query sequences. In each iteration, it computes an (*m*+*q*) × (*m*+*q*) distance matrix on sequences from both the backbone and queries using shared 6-mers. Then, it computes a guide tree using the distance matrix and builds an alignment. More specifically, for each node in the guide tree, MAFFT-add does an alignment computation only if a query sequence is involved at the node (i.e., at least one child has some query sequences). Otherwise, it simply uses the alignment from the backbone. There are other variants of MAFFT-add that use more accurate distance calculation and can result in improved alignment accuracy (at the cost of scalability). One of the most accurate variants is MAFFT-linsi-add which we briefly describe below.

### 2.4 MAFFT-linsi-add

The procedure to add sequences using MAFFT-linsi-add has two differences to MAFFT-add. First, MAFFT-linsi-add uses *localpair* (local pairwise alignment scores) for the distance matrix calculation, which is more accurate than shared 6-mers. Second, it only runs for one iteration of progressive alignment and uses at most 1000 iterations of iterative refinement after the progressive alignment finishes. MAFFT-add and MAFFT-linsi-add have runtimes that are at least quadratic in the input size, due to the *O*((*m*+*q*)^2^) distance matrix calculation, where there are q query sequences and m sequences in the provided backbone alignment. MAFFT-linsi-add is even less scalable since its distance calculation is more costly. In addition, MAFFT-linsi-add does many steps of refinement that further impact the runtime. Hence, the designers recommend that MAFFT-linsi-add be limited to at most a few hundred sequences [10].

## 3 New Methods: SALMA-add and SALMA

We have developed a technique to enable MAFFT-linsi-add to scale to ultra-large datasets. This new sequence-adding method is part of a two-stage MSA method, SALMA, that we present, and is of interest in its own right. Hence, we refer to this method as SALMA-add, and study it in comparison to UPP-add and WITCH-add.

The core idea of SALMA-add is to break the sequence-adding problem into multiple smaller sub-problems, on which MAFFT-linsi-add can efficiently and accurately run (i.e., divide-and-conquer). Specifically, instead of trying to run MAFFT-linsi-add on the full backbone alignment, we break the backbone alignment into smaller sub-alignments (modifying techniques from previous divide-and-conquer MSA estimation methods, PASTA [13] and MAGUS [25]). Then, for each query sequence, we use a statistical test (described below) to pick the best subset for the query sequence, and we run MAFFT-linsi-add to add the query sequence into the selected sub-alignment. Since the sub-alignments are all consistent with the parent backbone alignment, this enables us to add the query sequence into the backbone alignment.

We bound the sub-alignment sizes to at least 50 sequences and at most *u* sequences (where *u* is set by the user) in order to improve accuracy but also enable MAFFT-linsi-add to be run. The upper bound of u sequences is based on the recommendation in [10], and the lower bound on the subset size is based on observations that dense taxonomic sampling can improve alignment accuracy (thus, sub-alignments that are too small can reduce accuracy).

Based on the motivations above, the SALMA-add pipeline has one free parameter, *u*, and is the following:

1. Compute a backbone tree and build an eHMM as done by UPP-add, but instead of *A* = 10, use *A* = 50 as the decomposition stopping criterion.
2. Instead of using all HMMs/sub-alignments, use sub-alignments of sizes at most *u* (note that the result is that all sub-alignments have sizes between 50 and *u*).
3. Assign query sequences to the selected HMMs/sub-alignments the same way as UPP-add (i.e., best fitting HMM for each query based on bit-scores).
4. Use MAFFT-linsi-add to add assigned query sequences to each subalignment. If the number of sequences in the sub-alignment plus the number of assigned queries exceeds 500, then evenly divide the assigned queries into smaller subsets so that each sub-problem has at most 500 sequences.

Note that this design achieves two properties: all sub-problems on which we run MAFFT-linsi-add have at most 500 sequences in total (including both the backbone sub-alignment and the query sequences) and the query sequences are assigned to a sub-problem for which they are likely to be closely related (based on the bit-score calculation). These two properties together make for sub-problems that are small enough for MAFFT-linsi-add to run well on and closely related enough to have good accuracy.

SALMA addresses the problem of *de novo* alignment of datasets with sequence length heterogeneity. Like UPP and WITCH, SALMA has two stages: it first selects and aligns a subset of the input sequences, thus forming the backbone alignment, and then adds the remaining sequences into the backbone. For the first stage, SALMA is identical to UPP and WITCH, but for the second stage it uses SALMA-add.

## 4 Experimental design

### 4.1 Overview

Experiment 1 is used for the design phase of SALMA, where in Experiment 1(a) we evaluate the scalability of MAFFT-linsi-add and in Experiment 1(b) we tune the parameter *u* for SALMA (i.e., the upper bound for sizes of sub-alignments); these analyses are performed on training datasets only. Experiments 2 and 3 evaluate SALMA and SALMA-add (the step in SALMA that adds sequences into provided backbone alignments), respectively, in comparison to other methods on the testing datasets (disjoint from the training datasets). All methods are evaluated for alignment accuracy and runtime. All analyses were run on the UIUC Campus Cluster with 16 cores and 64 GB memory except for Rec and Res datasets (the largest two datasets we align), which we set the memory to 128 GB. The runtime limit is set to 12 hours for Experiment 1(a), 48 hours for analyses of Rec and Res, and 24 hours for the remaining experiments. See the supplementary materials for exact commands of all methods and for additional details.

### 4.2 Datasets

We use both simulated and biological datasets of nucleotide and protein to evaluate SALMA and SALMA-add. We focus on model conditions that exhibit sequence length heterogeneity (e.g., see Figure 1), with histograms for all datasets provided in supplementary materials Figures S4-S8. Empirical statistics for these datasets can be found in Table 1, where a higher p-distance (i.e., normalized Hamming distance, see supplementary materials for the definition) means higher substitution rates.

**Figure 1:**
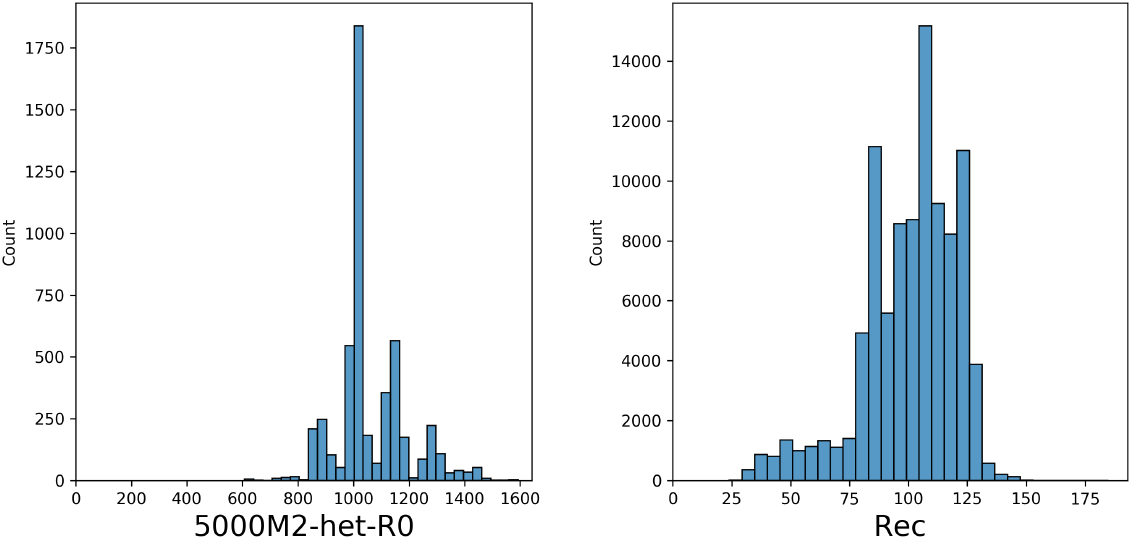
Sequence length histograms of replicate 0 of 5000M2-het (left) and Recombinase (Rec) (right).

**Table 1:**
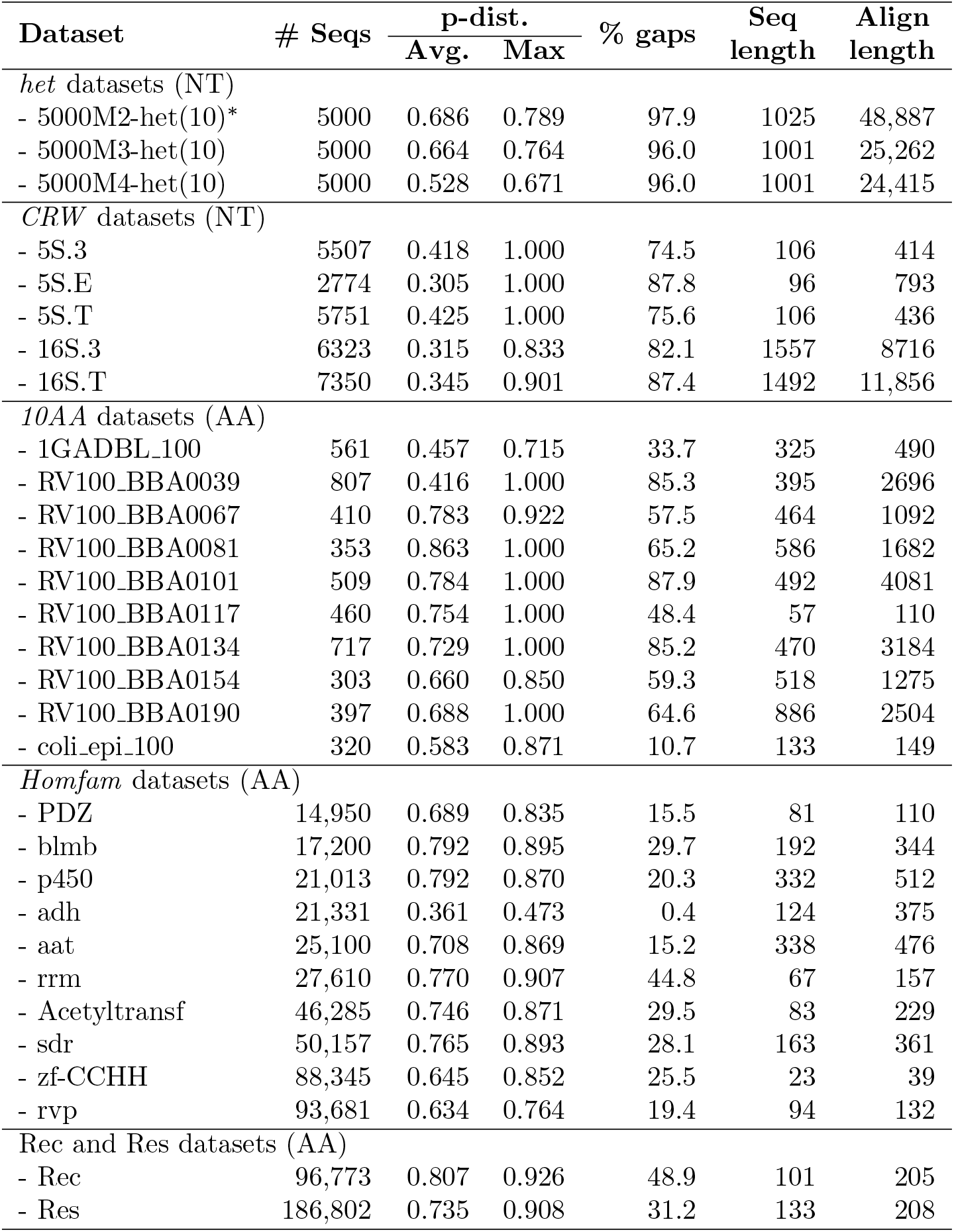
Empirical statistics of nucleotide (NT) and protein (AA) datasets. Datasets marked with (*) denote training datasets. The number of replicates (if more than one) is specified with a parenthesis after the dataset name. p-dist.: p-distance (i.e., normalized Hamming distance).

#### The het simulated datasets

We generated three new simulated conditions with evolutionary sequence length heterogeneity resulting from only from insertions and deletions during their evolution down model trees, rather than fragmentation. Each replicate has 5000 sequences and range in evolutionary rates (and hence alignment difficulty), with model condition 5000M2-het having the highest rate of evolution, 5000M3-het somewhat slower, and 5000M4-het the slowest. The model for sequence length heterogeneity assumes that insertion/deletion (“indel”) events, with a small probability, can be promoted to long indel events modeling infrequent large gain or loss of genic regions during the evolutionary process (e.g., domain-level indels). We used non-ultrametric model trees and simulated sequence evolution using INDELible [7]. The sequence length distribution for one replicate from these simulations is presented in Figure 1.

#### CRW

We include five large datasets from the Comparative Ribosomal Website [2]: 5S.3, 5S.E, 5S.T, 16S.3, and 16S.T, with 4507, 1773, 4751, 5323, and 6350 sequences, respectively. These datasets have been used in previous studies [3, 17, 22], exhibit sequence length heterogeneity, and have reference alignments based on secondary structures. We use the cleaned versions from [22], for which any ambiguity codes or entirely gapped columns are removed. We refer to these five datasets as *CRW*.

#### Homfam

We include the ten largest Homfam protein datasets [1, 24], which range from 14,950 to 93,681 sequences. Each of these datasets has a reference alignment on only a few sequences, and thus the alignment accuracy is only evaluated on a subset of sequences for each dataset.

#### 10AA

We include the “10AA” datasets that were used in prior studies to evaluate alignment methods [13, 17, 3]. These 10AA datasets are curated protein alignments based on structure that range from 320 to 807 sequences.

#### Serine Recombinases (Rec and Res)

We use datasets for two domains from serine recombinase. Protein sequences were taken from 350,378 GenBank bacterial and archaeal genome assemblies (supplementary materials) using Prodigal [8]. Serine recombinases were identified using the Pfam HMMs [15] Resolvase (Res), for the catalytic domain, and Recombinase (Rec), for the integrase-specific domain, using *hmmsearch* from the HMMER package with the “trusted cutoffs” supplied by Pfam. Standards phiC31 and Bxb1 were added. From the 199,090 unique protein sequences, the Rec and Res domains were separately extracted, using the boundaries determined by the HMM hits. As with the Homfam datasets, each of Rec and Res has a reference alignment on only a few of the sequences (i.e., the seed alignment from Pfam), and so alignment accuracy is evaluated only on these specific sequences.

#### Training vs. testing data

The het simulation has three model conditions; one was used in Experiment 1 as training data, and the other two were used in Experiments 2 and 3. All other datasets were used only for testing (i.e., in Experiments 2 and 3).

### 4.3 Other MSA methods in the study

We compare SALMA to Clustal-Omega [23], FAMSA [4], MAFFT [12], MAFFT-sparsecore [12], MAGUS [25], MUSCLE [5], PASTA [13], UPP (using a MAGUS backbone) [22], and WITCH [21]. We compare SALMA-add, the sequence-adding stage of SALMA, to the following sequence-adding methods: MAFFT-add (run in default mode), UPP-add, and WITCH-add. We omit MAFFT-linsi-add due to its scalability issues, as we establish in Experiment 1. MAFFT-sparsecore does not enable the user to provide the backbone alignment, and so is not included.

### 4.4 Backbone alignment and query sequences

We follow the UPP procedure to split each sequence dataset into a backbone set and a query set. For each dataset, the backbone set draws a subset of sequences from all full-length sequences (i.e., lengths 25% around the median length), while the query set contains all remaining sequences. Based on results from UPP2 [18], a large backbone set usually leads to better alignment accuracy on the final alignment of all sequences compared to using fewer full-length sequences in the backbone at the cost of runtime. Hence, we adopt the same backbone selection strategy by using all available full-length sequences.

We compute an alignment on the backbone set using MAGUS [25], because it provides improved accuracy compared to PASTA, the prior technique used in UPP [22]. However, if desired, the backbone alignment can be performed by any other preferred alignment method.

### 4.5 Evaluation metrics

We evaluate alignment methods with respect to wall-clock running time and alignment error. For alignment error, we compare estimated and reference alignments, reporting SPFN, SPFP, and the average of these two, defined as follows. Homologies are pairs of letters (nucleotides or amino acids), one from each of two sequences, that appear in the same column within the true or estimated alignment. The sum-of-pairs falsenegative (SPFN) rate is the fraction of pairs of homologies in the reference alignment but missing in the estimated alignment. The sum-of-pairs falsepositive (SPFP) rate is the fraction of pairs of homologies that are in the estimated alignment but missing in the reference alignment. These error rates are calculated using FastSP [14]. To evaluate alignment error for sequence-adding methods, we provide the same backbone alignment to each method and then evaluate the alignments induced on the added query sequences.

## 5 Results

### 5.1 Experiment 1: Designing SALMA

Experiment 1 involves two sub-experiments: 1(a) is for evaluating different ways of running MAFFT–add (either with–linsi or in default mode) for accuracy and scalability, and 1(b) is for designing the divide-and-conquer strategy in SALMA.

#### 5.1.1 Experiment 1(a) - scalability of MAFFT-linsi-add

In this experiment, we explore what sizes of datasets MAFFT-linsi-add can align without crashing or encountering memory issues. We use our training dataset 5000M2-het as the benchmark and either MAFFT-add or MAFFT-linsi-add to add 100, 200, 1000, or 2000 query sequences to the backbone.

Figure S1 (supplementary materials) shows the results for runtime and alignment error comparing MAFFT-add in default mode (which does not use MAFFT-linsi-add except on small datasets) and MAFFT-linsi-add, when adding sequences into large backbone alignments. MAFFT-linsi-add encountered out-of-memory issues on several replicates, and in some cases did not complete within 12 hours. However, when MAFFT-linsi-add was able to run, it was more accurate (sometimes substantially more accurate) than MAFFT-add run in default mode, but used more time. These results show the advantage in using MAFFT-linsi-add, but also its limitations to small datasets.

#### 5.1.2 Experiment 1(b): Tuning parameter *u*

The parameter *u* specifies the lower bound on the size of sub-alignments we use for MAFFT-linsi-add in SALMA. We explore the impact of this parameter by varying *u* between 100, 200, and 400, using the training dataset 5000M2-het. We consider two ways of defining the backbone sequences: all full-length sequences (i.e., the UPP2 strategy) and at most 1000 full-length sequences (i.e., the UPP strategy). Both strategies add the remaining sequences to the same backbone alignment using SALMA-add.

Results for SALMA with different settings of *u* on all sequences of 5000M2-het (10 replicates) are presented in the supplementary materials (Figure S2). For a fixed backbone size, the differences between SALMA variants are negligible for both SPFN and SPFP. Using all full-length sequences for the backbone has a lower SPFP than using just 1000 full-length sequences, while both have similar SPFN. Overall, the “all” backbone strategy has a lower alignment error than the “1000” backbone for all SALMA variants. For runtime, using “1000” backbone is slightly faster than “all” backbone, while SALMA with smaller *u* (i.e., *u* = 100) is faster.

Based on the training results, we choose *u* = 100 and use the “all” backbone selection strategy for SALMA in the following experiments, because this variant gives the best alignment accuracy while having a relatively lower running time than other variants.

### 5.2 Experiment 2: Evaluating SALMA

We compare SALMA to leading MSA methods that can run on large datasets. Comparisons to MAFFT-sparsecore [12], FAMSA [4], MAGUS [25], and WITCH [21] are provided here, but comparisons to the remaining methods (i.e., Clustal-Omega [23], MUSCLE [5], PASTA [13], MAFFT [12], and MAGUS+UPP (i.e., UPP with MAGUS backbone) [22]) are summarized here and provided in the supplementary materials, Tables S3 and S4.

Because of the large size for the Rec and Res datasets, we modify the setting for *A* (which specifies the stopping rule for the decomposition) within SALMA and WITCH: we use *A* = 50 instead of the default *A* = 10. For the Res dataset, which is the larger of the two datasets, we use MAGUS in recursive mode to compute an alignment on the full dataset. When used within two-stage methods (e.g., SALMA and WITCH), we also use MAGUS in recursive mode for the backbone alignment, because it otherwise encountered out-of-memory issues when given 128 GB.

We present all benchmark results in Table 2. We take SPFN and SPFP of each method over all replicates within a collection of datasets (e.g., *CRW* or *Homfam*), and calculate their averages. We boldface the best methods for each benchmark and break ties with a margin of 0.01 (e.g., a SPFN of 0.110 and 0.119 are both considered the best).

**Table 2:**
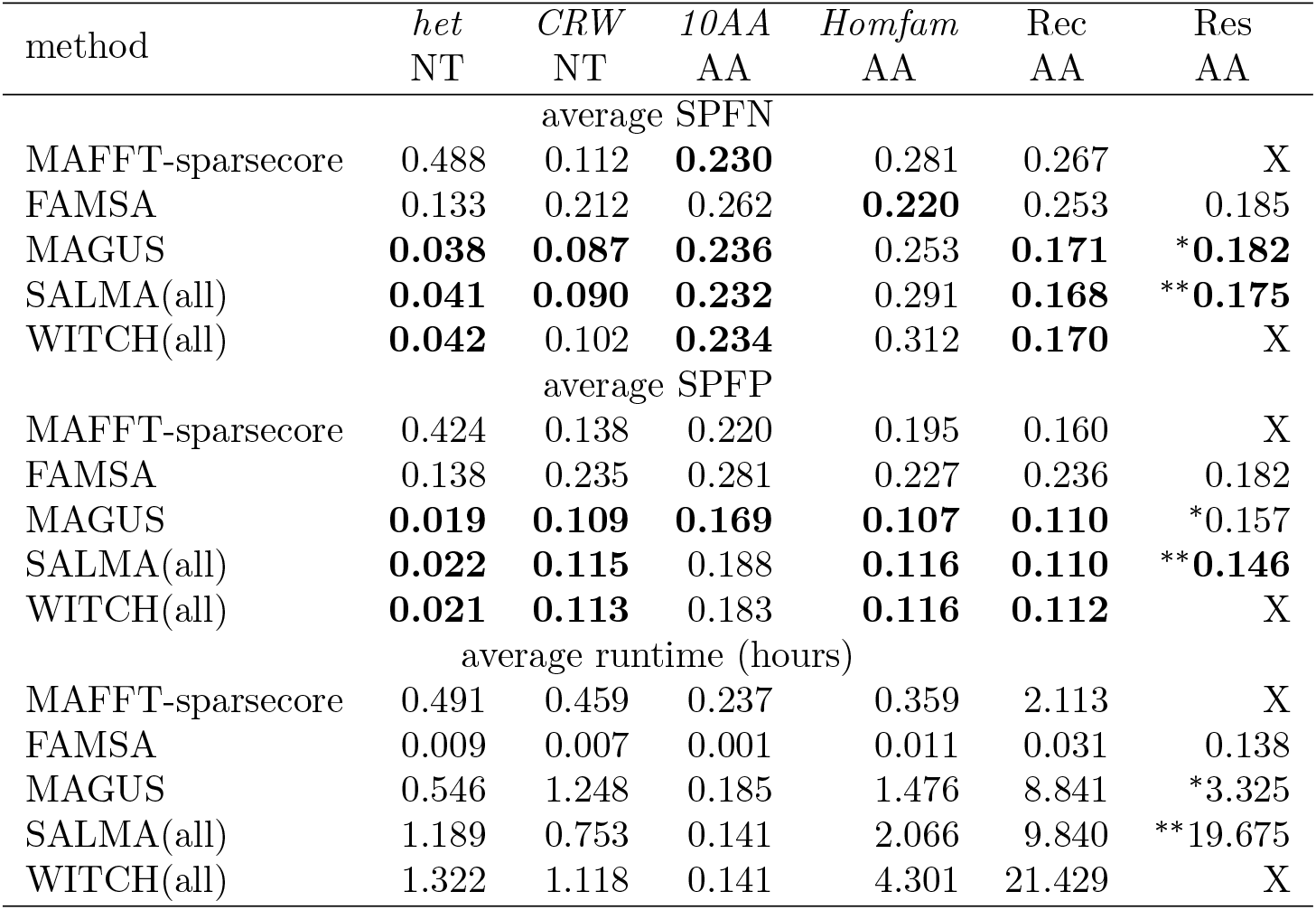
Experiment 2: Alignment error and runtime (in hours) over several nucleotide (NT) and protein (AA) benchmarks for different alignment methods. The notation “(all)” for SALMA and WITCH denotes that they use all fulllength sequences to construct the backbone alignment and add the remaining sequences. We boldface the methods that are tied for best (i.e., within 0.01 of the best) for each benchmark. A cross mark (“X”) means that the method either encountered out-of-memory issues during the run or did not finish within the time limit. * MAGUS was run in recursive mode on the Res dataset. **The MAGUS backbone alignment was run in recursive mode on Res.

The first two columns of Table 2 show alignment results on two sets of nucleotide benchmarks, *het* and CRW. *het* contains both 5000M3-het and 5000M4-het (10 replicates each). MAGUS, WITCH, and SALMA are the best methods for both SPFN and SPFP. FAMSA is also accurate but not as accurate as the three methods mentioned earlier, while MAFFT-sparsecore has both high SPFN and SPFP. For the five datasets from *CRW*, SALMA ties with MAGUS to be the most accurate method for both SPFN and SPFP, while WITCH has a similar SPFP but slightly higher SPFN. MAFFT-sparsecore is slightly worse than the best methods for both SPFN and SPFP, while FAMSA is the least accurate.

The last four columns of Table 2 show alignment results on a collection of protein benchmarks: *10AA* (10 datasets), *Homfam* (the 10 largest Homfam datasets), Rec, and Res. For 10AA, all methods except FAMSA have similarly low SPFN. SPFP shows a larger gap between methods. MAGUS has the lowest SPFP, and SALMA and WITCH are the second best. FAMSA has noticeably higher SPFP than the remaining methods. For *Homfam*, FAMSA is the best method in terms of SPFN. MAGUS comes as the second best and is followed by MAFFT-sparsecore. Both SALMA and WITCH have high SPFN, with SALMA being slightly better. On the other hand, SALMA, WITCH, and MAGUS are the best in SPFP, and FAMSA has a much higher SPFP than the remaining methods. For Rec and Res datasets, MAFFT-sparsecore failed to finish on Res due to out-of-memory issues (128 GB limit), and WITCH failed to finish on Res due to time limit (48 hours). SALMA is the best method in terms of SPFN and SPFP for both datasets. MAGUS and WITCH also are the best for Rec, and MAGUS comes as the second for the Res dataset.

FAMSA is the fastest method and is so by a large margin against the remaining methods. SALMA is among the slowest methods for most datasets, except the smallest ones (i.e., *CRW* and *10AA*). However, SALMA can scale to the largest dataset (Res) without running out of memory. In contrast, WITCH and MAFFT-sparsecore both fail to complete on the Res dataset due to memory issues.

Results for the other methods we evaluated (i.e., Clustal-Omega, MUSCLE, default MAFFT, PASTA, and UPP using a MAGUS backbone) are presented in Tables S3 and S4 (supplementary materials) and summarized here. MUSCLE failed to run on large datasets (e.g., two largest Homfam datasets and both Rec and Res). PASTA is the most accurate on *het* in terms of both SPFN and SPFP, but is otherwise strictly less accurate and slower than MAGUS. UPP using a MAGUS backbone has similar accuracy compared to WITCH. Thus, of the full set of methods we considered, none of these other methods needs further consideration.

### 5.3 Experiment 3: Evaluating SALMA-add

In this experiment, we use the same benchmarks from Experiment 2 and directly compare the alignment accuracy for adding sequences to existing backbone alignments using UPP-add, WITCH-add, SALMA-add, and MAFFT-add. Four of the Homfam datasets and Res have fewer than two query sequences with reference alignments and thus are omitted.

Figure 2 shows the SPFN and runtime in hours of methods for adding sequences using the same datasets from Experiment 2, where backbones have all full-length sequences and are aligned with MAGUS. In terms of SPFN, SALMA-add is the most accurate sequence-adding method except on *Homfam*, on which MAFFT-add has the lowest SPFN. For all benchmarks, WITCH-add is consistently more accurate than UPP-add for SPFN, MAFFT-add also has comparable SPFN to WITCH-add and UPP-add, but it has noticeably higher SPFN on the *het* benchmark.

**Figure 2:**
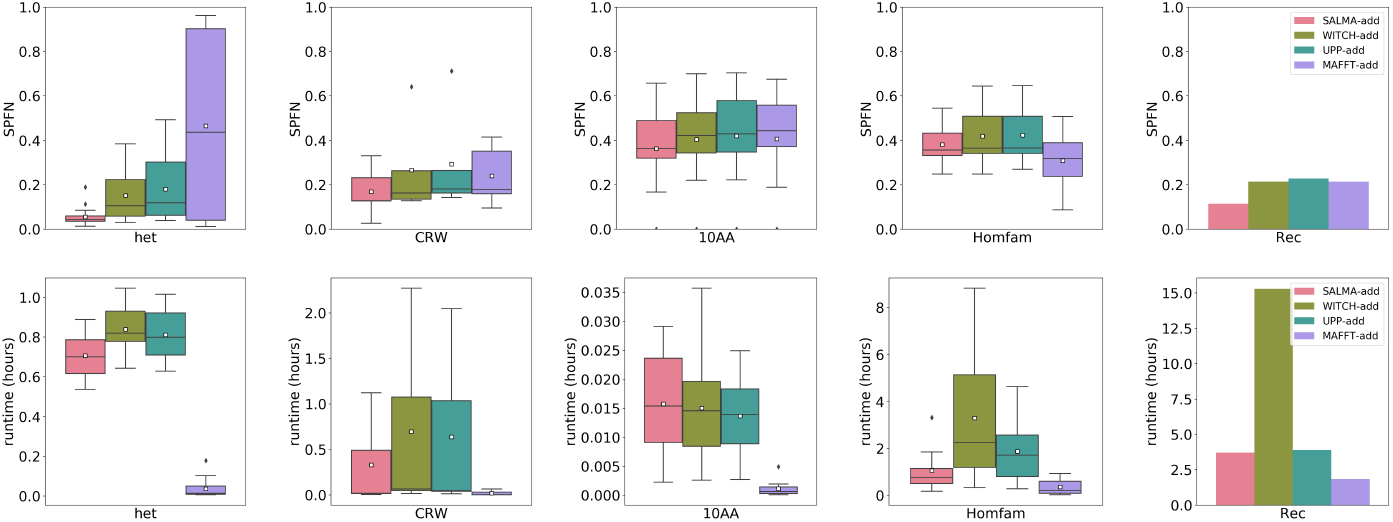
Experiment 3: SPFN (top 5 panels) and runtime in hours (bottom 5 panels) of four methods for adding sequences to existing backbone alignments. Backbone alignments are the same for all methods and are constructed by aligning all full-length sequences with MAGUS, and SPFN is evaluated on the added sequences only. SPFP can be found in the supplementary materials (Figure S3). From left to right: *het* consists of 5000M3-het and 5000M4-het, each with 10 replicates, *CRW* consists of five selected CRW datasets, *10AA* consists of 10 biological protein datasets, *Homfam* consists of 6 out of the largest 10 Homfam datasets (four of them have fewer than two query sequences with reference alignments and are omitted), and the last one is Rec. Res is not shown because it has fewer than two query sequences with reference alignment.

In terms of SPFP (see Supplementary Figure S3), SALMA-add is also usually the best except on *10AA*, on which UPP-add has the lowest SPFP. MAFFT-add often has the highest SPFP.

For runtime, MAFFT-add is the fastest sequence-adding method. SALMA-add is usually faster than both UPP-add and WITCH-add except for 10AA. All methods are fast on 10AA (finished within 3 minutes), and SALMA-add is slightly slower than UPP-add and WITCH-add. WITCHadd is the slowest sequence-adding method among the four, particularly on large datasets such as *Homfam* and Rec.

## 6 Discussion

Our study showed that SALMA provides accuracy advantages over other methods on a wide range of nucleotide and protein benchmarks. Furthermore, SALMA-add, the step in SALMA that performs sequence addition, provides superior accuracy for adding sequences into existing backbone alignments compared to UPP-add and WITCH-add, and is able to scale to much larger datasets than MAFFT-linsi-add.

It is easy to see why SALMA can scale to large datasets, as its design ensures that no problem is ever too big for MAFFT-linsi-add to complete. SALMA’s accuracy depends on the two-stage approach. Since SALMA differs from UPP and WITCH only in the second stage, understanding why SALMA is more accurate than UPP and WITCH amounts to understanding why SALMA-add is more accurate than UPP-add and WITCH-add. Both UPP-add and WITCH-add use eHMMs to infer homologies, while SALMA-add relies on MAFFT-linsi-add (albeit in a divide-and-conquer framework). This clearly indicates that MAFFT-linsi-add provides some advantage over sequence-adding techniques that use eHMMs.

It is also worth noting that where SALMA really excels is in its high recall (i.e., its ability to detect homologies). When compared to other sequence-adding methods, SALMA has the lowest SPFN (highest recall) on all benchmarks in this study, except for the Homfam benchmark, and most of the time the improvement on other methods is substantial. We also note that SALMA has generally low SPFP scores, although UPP-add and WITCH-add sometimes are better.

## 7 Conclusions

This study presented SALMA, a new multiple sequence alignment method specifically designed to align datasets with sequence length heterogeneity. The innovative part of SALMA is its second stage, SALMA-add, which adds query sequences into a backbone alignment computed in the first stage. SALMA-add uses a divide-and-conquer approach to enable MAFFT-linsi-add to scale to large datasets with very high accuracy. Both SALMA and SALMA-add provide accuracy advantages over other methods when aligning datasets with sequence length heterogeneity, as our study shows, and can scale to very large datasets.

This study suggests the potential for even further improvements in methods for aligning datasets that exhibit substantial sequence length heterogeneity, which is a problem that is of increasing importance in modern sequence dataset generation and analysis.

## Supporting information

"See the supplementary materials"

## 8 Acknowledgements

This work was supported by the US National Science Foundation grant 2006069 (to TW). It was also supported by the U.S. Department of Energy, Office of Science, Office of Biological and Environmental Research under the Secure Biosystems Design Initiative and by the Laboratory Directed Research and Development (LDRD) program of Sandia National Laboratories, which is a multimission laboratory managed and operated by National Technology and Engineering Solutions of Sandia, LLC, a wholly owned subsidiary of Honeywell International Inc, for the U.S. Department of Energy’s National Nuclear Security Administration under contract DE-NA0003525.

