## Supplementary material for "SALMA: Scalable ALignment using MAFFT-Add": "See the supplementary materials"

**S5 Experiment 1(a): MAFFT-linsi-add scalability issue on 5000-taxon datasets** 8

**S6 Additional Definitions** 8

**S7 Additional Figures** 9

**S8 Additional Tables** 15

### S1 Commands for software

All commands are based on the assumption that 16 CPU cores are available.

#### S1.1 Backbone generation and sequence-adding methods

1. MAGUS backbone alignment (GitHub version committed on April 5th 2021):

```
$ python3 magus.py --recurse false -np 16  \
    -i [unaligned backbone sequences] -d [outdir] \
    -o [output alignment]
```

2. FastTree2 backbone tree (`-nt` for nucleotide or none for amino acids, v2.1 multi-threaded version):

```
$ FastTreeMP {/-nt} -gtr [backbone alignment] > [backbone tree
    ]
```

3. UPP-add (UPP v4.5.0):

```
$ python3 run_upp.py -x 16 -s [query sequences path] \
    -a [backbone alignment] -t [backbone tree] -p [tempdir] \
    -d [outdir] -o [job name] --molecule [dna/amino] \
    -A [decomposition size, default 10]
```

4. WITCH-add (Github commit head **a5fff8b**):

```
$ python3 witch.py -t 16 -q [query sequences path] \
    -b [backbone alignment] -e [backbone tree] -d [outdir] \
    -o [output name] -A [decomposition size, default 10] \
    --molecule [dna/amino]
```

5. MAFFT/MAFFT-linsi --add (MAFFT v7.490):

```
$ [mafft/mafft-linsi] --quiet --thread 16 --add [query
    sequences] \
    [backbone alignment] > [output alignment]
```

#### S1.2 Standard alignment methods

1. MAGUS (GitHub version committed on April 5th 2021):

```
$ python3 magus.py --recurse {true/false} -np 16  \
    -i [unaligned sequences] -d [outdir] \
    -o [output alignment]
```

2. FAMSA (v2.0.3):

```
$ famsa -t 16 [unaligned sequences] [output alignment]
```

3. MAFFT-sparsecore (MAFFT v7.490; also require Ruby interpreter installed):

```
$ mafft-sparsecore.rb -C ‘--quiet --thread 16’ \
    -A ‘--quiet --thread 16’ -i [unaligned sequences] \
    > [output alignment]
```

4. PASTA (v1.9.0):

```
$ python3 run_pasta.py -i [unaligned sequences] -o [outdir] \
    --temporaries [tempdir] --keeptemp --num-cpus=16
```

5. MAFFT (default, v7.490):

```
$ mafft --quiet --thread 16 [unaligned sequences] \
    > [output alignment]
```

6. MUSCLE (v3.8.31):

```
$ muscle -maxiters 2 -in [sequence file] \
    -out [output alignment]
```

7. Clustal-Omega (v1.2.4):

```
$ clustalo --infile=[unaligned sequences] --infmt=fasta \
    --outfile=[output alignment] --outfmt=fasta \
    --threads=16
```

##### S1.3 Evaluations

1. FastSP (v1.7.1) to obtain SPFN and SPFP. Alignment error is then obtained by averaging SPFN and SPFP. The "-ml -mlr" options will ignore columns with lowercase letters (considered as insertions in UPP and WITCH):

```
$ java -jar FastSP.jar -ml -mlr -e [estimated alignment] \
    -r [reference alignment] -o [output file]
```

2. Runtime is obtained by adding the following command before any software:

```
$ { /usr/bin/time -v [software command and options] ; } \
    > [runtime information]
```

3. Empirical dataset statistics (i.e., p-distance, % gaps, etc.) are obtained using the following script (`https://wiki.illinois.edu/wiki/display/warnowlab/Get+alignment+statistics`). Alternatively link from Dropbox (`https://www.dropbox.com/s/wqt8hpvqkdjge04/alignment_stats.zip?dl=0`):

```
$ java -cp ./alignment_stat AlignmentStatistics \
    [alignment file] [output file]
```

#### S2 Data Availability

**het.** The newly generated datasets, 5000M2-het, 5000M3-het, and 5000M4-het, with evolutionary sequence length heterogeneity, are available at `https://databank.illinois.edu/datasets/IDB-3974819`.

**CRW.** The cleaned version of the five CRW datasets (5S.3, 5S.E, 5S.T, 16S.3, and 16S.T) are available at `https://databank.illinois.edu/datasets/IDB-2419626`.

**10AA.** 10AA datasets are available at https://drive.google.com/file/d/0B0lcoFFOYQf8d0IzdkJfdG9jUEk/view?resourcekey=0-HLUfGH32UNi8cDgwRt27wA.

**Homfam.** Homfam datasets are available at https://drive.google.com/file/d/0B0lcoFFOYQf8Z1NDMERacUN4LTA/view?resourcekey=0-SPM-574k-8_T7auVfFb__w. All 20 datasets are available, and we only use the largest 10 in this study. For each Homfam dataset, there are only a few sequences with reference alignments. The actual numbers of sequences with reference alignments are:

- Acetyltransf: 6
- PDZ: 6
- aat: 10
- adh: 5
- blmb: 6
- p450: 12
- rrm: 20
- rvp: 6
- sdr: 13
- zf-CCHH: 15

**Rec and Res.** Locations for the Rec and Res datasets are provided at https://tandy.cs.illinois.edu/datasets.html. Pfam seed sequences are included for both datasets (66 and 110 sequences for Rec and Res, respectively).

**Backbone and query split.** For SALMA, WITCH, and MAGUS+UPP, the backbone/query sequences, as well as their reference alignments of each dataset, are available at https://tandy.cs.illinois.edu/datasets.html. All full-length sequences are included in the backbone set, while the remaining sequences form the query set for each dataset. All sequences and their reference alignments of each dataset are also included.

### S3 Dataset Generation: het

We present details for the generation of our new simulated conditions below. Our simulation parameters are based on those of the 1000M series of the ROSE simulated dataset [Liu et al., 2009]. We uploaded all of our INDELible control files generating this data to https://github.com/ThisBioLife/5000M-234-het to allow easy reproduction.

| **Condition** | $l$ **(tree scale)** | $r$ **(indel rate)** |
|---|---|---|
| 5000M2 | 30 | $4.4 \times 10^{-4}$ |
| 5000M3 | 20 | $3.3 \times 10^{-4}$ |
| 5000M4 | 5 | $1.32 \times 10^{-3}$ |

**Table S1: Parameters for generating the 5000M-het series data, where $l$ is the tree scale parameter, defining the maximum path-length in the non-ultrametric model tree, and $r$ defines the indel rate.**

#### S3.1 Model trees

Our model trees were generated by a two-step process. We first generated random 5000-species birth-death trees (one tree per replicate) using the DendroPy [Sukumaran and Holder, 2010] `treesim.birth_death_tree` function, setting the birth rate initially at 1 with the standard deviation of the change to the birth rate set to 0.2 (see the documentation on this Dendropy function on the birth-death process and the implication of this parameter), with a zero death rate. Then, taking only the topology of the tree generated in the prior step, we instructed INDELible to assign branch lengths to the tree such that the resulting tree is both non-ultrametric and has a maximum path-length $l$. We vary $l$ to control the scale of the tree and hence the rate of evolution of the condition. The choice of $l$ across conditions can be seen in Table S1.

#### S3.2 Indel length distribution & indel rates

Our model for sequence length heterogeneity assumes a base ("short indel") distribution of indel lengths. In our case, we simply took the "medium" length distribution from the "M"-series ROSE simulated datasets [Liu et al., 2009] as the short indel distribution. Given an indel event, with probability $p = 0.85$, it draws its length from the short indel distribution. Otherwise, it draws its length from a long indel distribution, in our case set to $\mathrm{NB}(130, 0.5)$. The indel length distribution is thus equivalently a mixture distribution of the short indel distribution (prior probability 0.85) and the long indel distribution (prior probability 0.15). We directly computed the probability mass function (PMF) of this mixture distribution, truncated the PMF, and fed the truncated PMF to INDELible as part of the input. The truncated PMF can be found alongside the uploaded control files in the Github repository linked.

Similar to the original ROSE dataset, we also varied the indel rates across conditions, with the choices for the indel rate $r$ shown in Table S1. The indel rates were chosen analogous to the original indel rates of the ROSE dataset.

#### S3.3 Other parameters

The rest of the parameters (e.g., GTR+$\Gamma$ parameters and the initial sequence length) were chosen to be the same as the ROSE simulated dataset generated for the SATé study [Liu et al., 2009] and can be found in the uploaded control files (`https://github.com/ThisBioLife/5000M-234-het`). We also provide the parameters here:

| GTR Parameters | |
|---|---|
| T->C | 1.2619573850882344 |
| T->A | 0.14005536945585983 |
| T->G | 0.2877830346145434 |
| C->A | 0.35766826674033914 |
| C->G | 0.3082674310184066 |
| A->G | 1 |

**Table S2: GTR matrix parameters for generating the 5000M-het series dataset. The parameters are set according to the parameters for the ROSE simulated data [Liu et al., 2009].**

- GTR parameters: see Table S2
- Stationary frequencies (`TCAG`): 0.311475, 0.191363, 0.300414, 0.196748
- $\alpha$ (for the gamma distribution): 1
- Initial sequence length: 1000

### S4 Dataset Generation: Rec and Res

#### S4.1 Background

There exist developed software that maps mobile DNAs within bacterial and archaeal genomes [Hudson et al., 2015, Mageeney et al., 2020], where each mapping is associated with the sequence of the integrase enzyme that catalyzes the site-specific integration of the mobile DNA.

#### S4.2 Generation

Protein sequences were taken from 350,378 bacterial and archaeal genome sequences (list available at TBD) using Prodigal [Hyatt et al., 2010]. Serine recombinases were identified using the Pfam HMMs [Mistry et al., 2021] Resolvase (Res), for the catalytic domain, and Recombinase (Rec), for the integrase-specific domain. Seed sequences are included for evaluation later (112 and 66 Pfam seed sequences for Res and Rec, respectively).

Link to the Res seed sequences: `http://pfam.xfam.org/family/PF00239#tabview=tab3`

Link to the Rec seed sequences: `http://pfam.xfam.org/family/PF07508#tabview=tab3`

**Sequence trimming** Two separate sequence datasets (i.e., Rec and Res) were prepared for the serine recombinases trimmed either according to the Recombinase or Resolvase HMM hits.

#### S4.3 Evaluation

We compare the estimated alignments to the seed Pfam alignments for both Rec and Res to obtain SPFN and SPFP (see main text for definitions).

#### S5 Experiment 1(a): MAFFT-linsi-add scalability issue on 5000-taxon datasets

Our runtime environment is 16 cores, 64 GB memory, and 12-hour runtime limit. We tried running MAFFT-linsi --add (MAFFT-linsi-add) on our training 5000M2 datasets, where for each replicate, we added 4000 query sequences to a 1000-taxon full-length backbone alignment. We encountered either out-of-memory issues (64 GB memory limit) or crashes. We also conducted an experiment on exploring the scalability of MAFFT-linsi-add by altering the number of queries (Experiment 1(a)).

The out-of-memory error message looks like the following:

```
slurmstepd: error: Detected 1 oom-kill event(s) in
StepId=5376434.batch cgroup. Some of your processes may have been
killed by the cgroup out-of-memory handler.
```

The crash error message looks like the following:

```
Command exited with non-zero status 1
```

#### S6 Additional Definitions

**Normalized Hamming distance.** The Hamming distance between two (aligned) sequences is defined as the number of columns that have different symbols (but not gaps).

Normalized Hamming distance is defined as the Hamming distance divided by the total number of columns that both sequences have symbols (but not gaps). It thus would be a number between 0.0 and 1.0, inclusively.

#### S7 Additional Figures

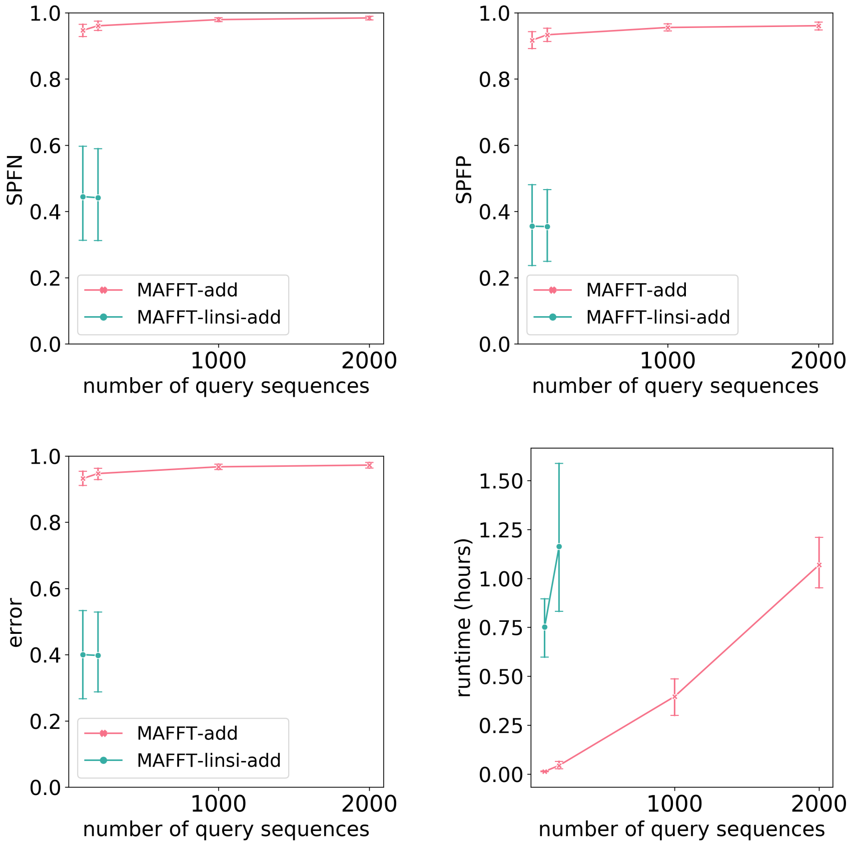

**Figure S1: Experiment 1(a): SPFN (top left), SPFP (top right), alignment error (average of SPFN and SPFP, bottom left), and runtime in hours (bottom right) of MAFFT-add and MAFFT-linsi-add for adding 100, 200, 1000, or 2000 sequences to a 1000-taxon backbone alignment. The dataset used is 5000M2-het with 10 replicates, where 1000 full-length sequences are randomly selected and aligned with MAGUS [Smirnov and Warnow, 2021] to form the backbone alignment. We exclude replicate 4 because MAFFT-linsi-add encountered out-of-memory issues when adding 100 or 200 query sequences. Additionally, MAFFT-linsi-add either encountered out-of-memory issues or did not complete within 12 hours when adding 1000 or 2000 query sequences and thus is not shown.**

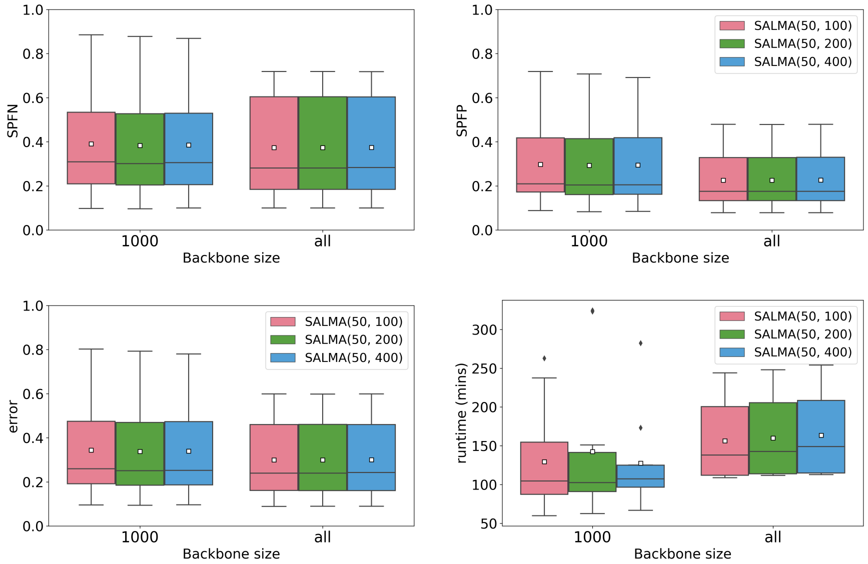

**Figure S2: Experiment 1(b): SPFN (top left), SPFP (top right), alignment error (average of SPFN and SPFP, bottom left), and runtime in hours (bottom right) of SALMA with various settings for adding query sequences (i.e., all remaining sequences) to a backbone alignment with either 1000 or all full-length sequences of 5000M2-het (10 replicates). Alignment metrics are calculated on all sequences, and white squares mark the averages. SALMA variants are marked as SALMA(50, $u$), for which $u = \{100, 200, 400\}$ controls the maximum size of sub-alignments to use for MAFFT-linsi-add. For example, $u = 100$ means that we only use sub-alignments with $[50, 100]$ sequences to add the query sequences.**

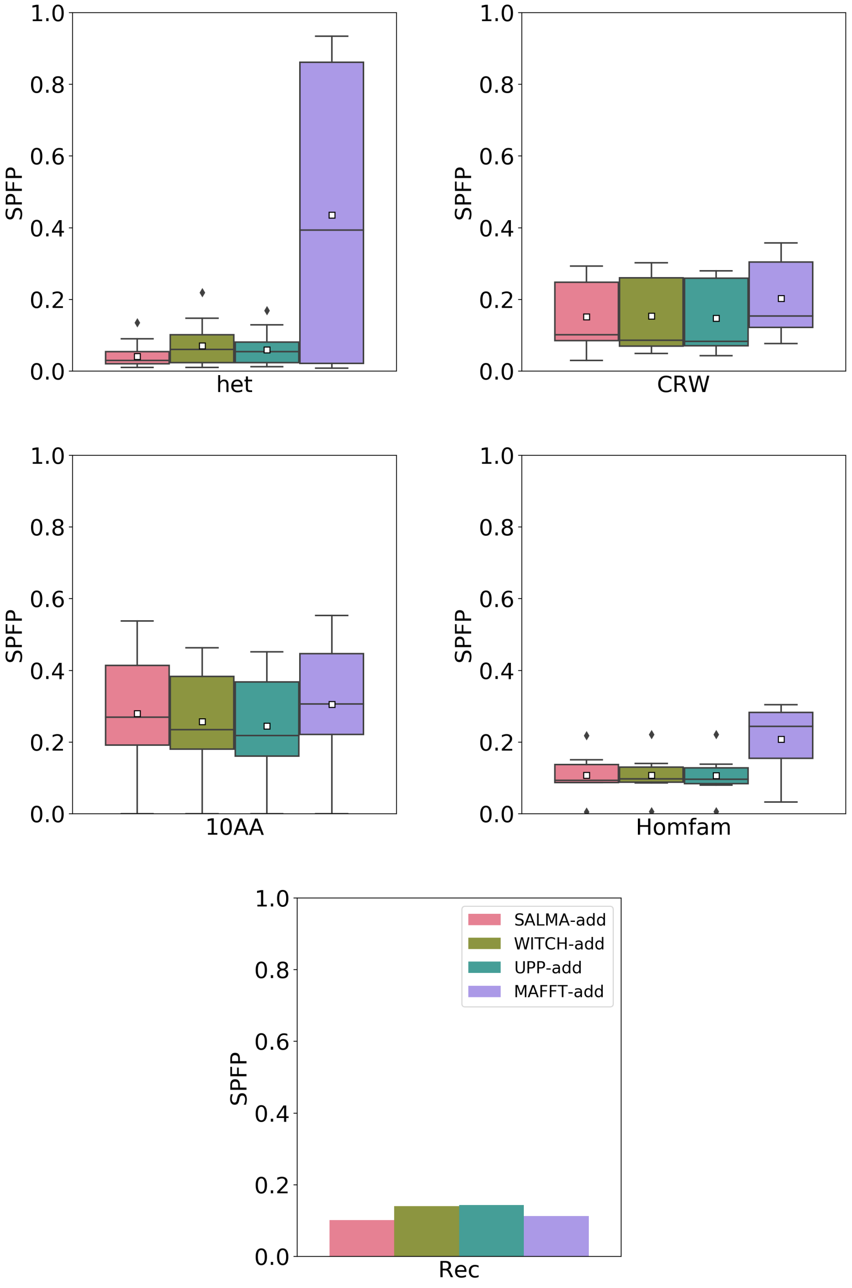

**Figure S3: Experiment 3: SPFP of four methods for adding sequences to existing backbone alignments. Backbone alignments are the same for all methods and are constructed by aligning all full-length sequences with MAGUS, and SPFP is evaluated on the added sequences only. From left to right, top to bottom: het consists of 5000M3-het and 5000M4-het, each with 10 replicates, CRW consists of five selected CRW datasets, 10AA consists of 10 simulated protein datasets, Homfam consists of 6 out of the largest 10 Homfam datasets (four of them have fewer than two query sequences with reference alignments and are omitted), and the last one is Rec. Res is not shown because it has fewer than two query sequences with reference alignment.**

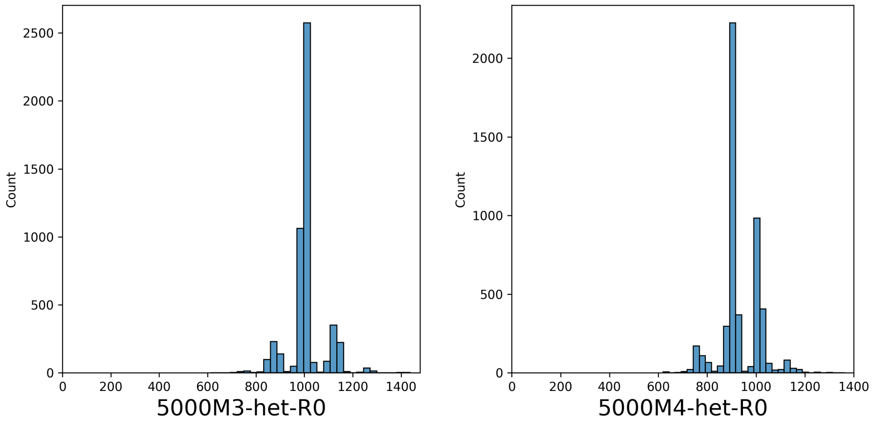

**Figure S4: Sequence length histogram of replicate 0 of 5000M3-het and 5000M4-het.**

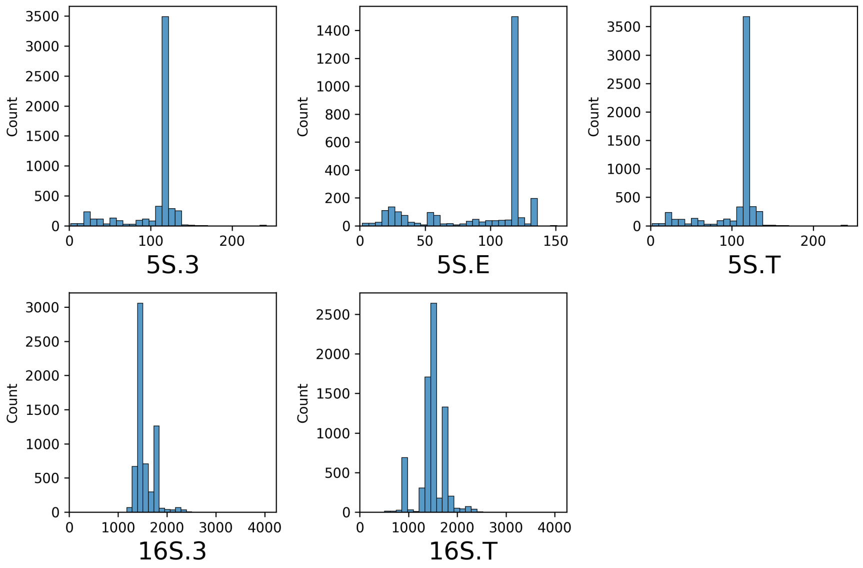

**Figure S5: Sequence length histograms of the five CRW datasets.**

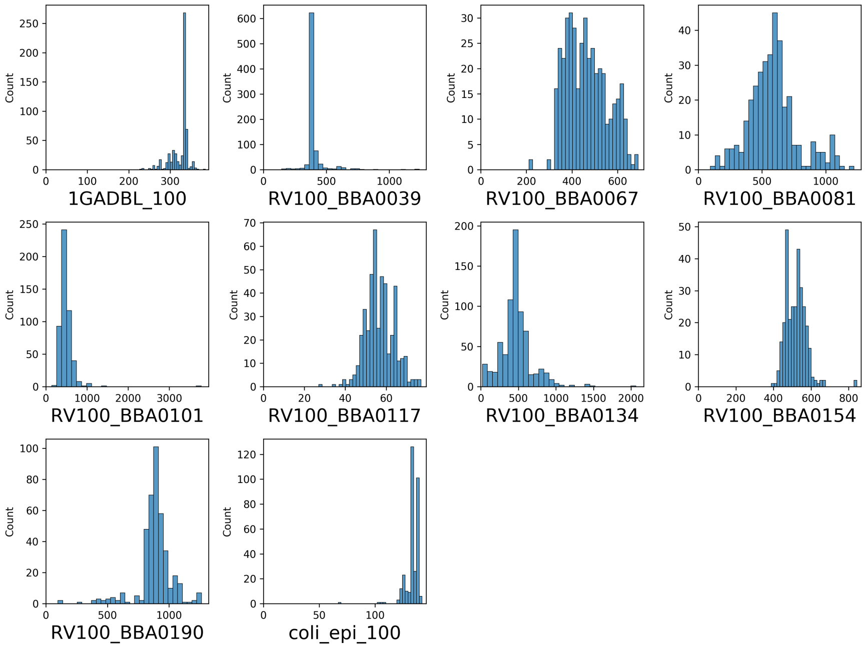

**Figure S6: Sequence length histograms of the 10AA simulated protein datasets.**

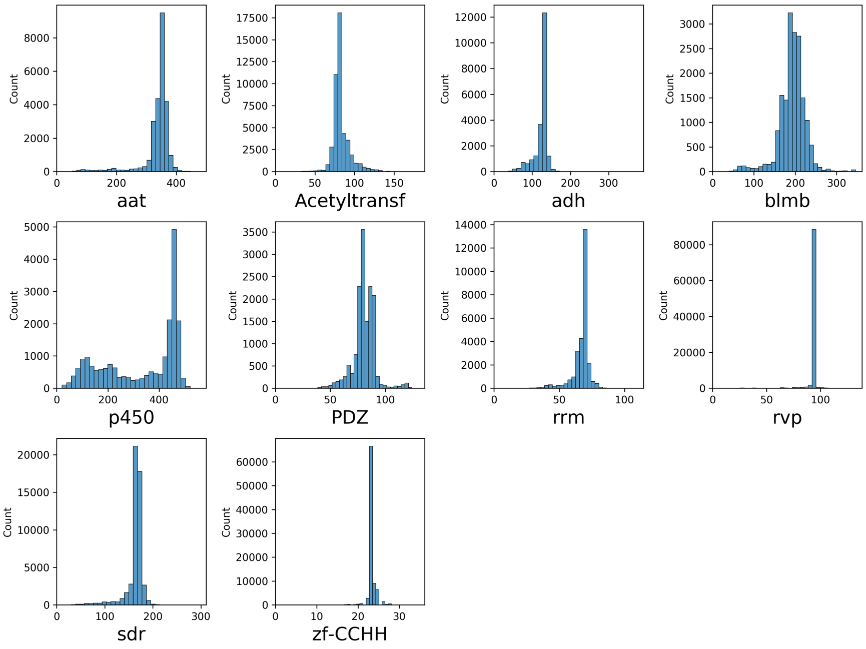

**Figure S7: Sequence length histograms of the 10 largest Homfam datasets.**

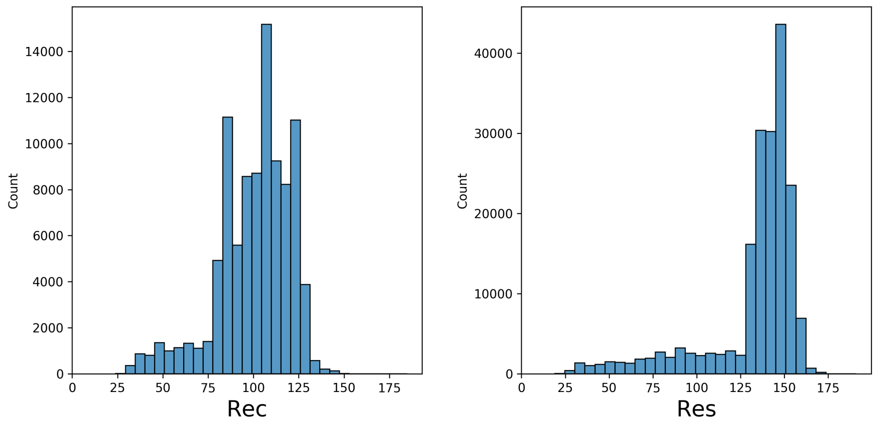

**Figure S8: Sequence length histograms of the Rec and Res datasets.**

#### S8 Additional Tables

| method | 5000M3-het | 5000M4-het |
|---|---|---|
| average SPFN | | |
| Clustal-Omega | 0.947 | 0.741 |
| MUSCLE | 0.974 | 0.273 |
| MAFFT(default) | 0.985 | 0.164 |
| PASTA | 0.045 | 0.016 |
| MAFFT-sparsecore | 0.922 | 0.054 |
| FAMSA | 0.208 | 0.059 |
| MAGUS | 0.046 | 0.030 |
| SALMA(all) | 0.054 | 0.029 |
| WITCH(all) | 0.055 | 0.030 |
| MAGUS+UPP(all) | 0.056 | 0.030 |
| average SPFP | | |
| Clustal-Omega | 0.749 | 0.469 |
| MUSCLE | 0.928 | 0.166 |
| MAFFT(default) | 0.962 | 0.091 |
| PASTA | 0.044 | 0.015 |
| MAFFT-sparsecore | 0.812 | 0.035 |
| FAMSA | 0.214 | 0.063 |
| MAGUS | 0.029 | 0.010 |
| SALMA(all) | 0.032 | 0.011 |
| WITCH(all) | 0.031 | 0.011 |
| MAGUS+UPP(all) | 0.030 | 0.011 |
| average runtime (hours) | | |
| Clustal-Omega | 1.496 | 0.257 |
| MUSCLE | 0.273 | 0.627 |
| MAFFT(default) | 0.036 | 0.015 |
| PASTA | 1.873 | 1.186 |
| MAFFT-sparsecore | 0.732 | 0.251 |
| FAMSA | 0.012 | 0.006 |
| MAGUS | 0.611 | 0.481 |
| SALMA(all) | 1.249 | 1.129 |
| WITCH(all) | 1.380 | 1.263 |
| MAGUS+UPP(all) | 1.335 | 1.253 |

**Table S3: Experiment 2: Average SPFN, SPFP, and runtime in hours for all tested methods on simulated datasets. 5000M3-het and 5000M4-het each has 10 replicates. "(all)" for SALMA, WITCH, and MAGUS+UPP denotes that they use all full-length sequences to form the backbone alignment.**

| method | *CRW* | *10AA* | *Homfam(8)* | *Homfam(10)* | Rec | Res |
|---|---|---|---|---|---|---|
| | | average SPFN | | | | |
| Clustal-Omega | 0.357 | 0.265 | 0.466 | 0.445 | 0.717 | 0.786 |
| MUSCLE | 0.623 | 0.298 | 0.698 | X | X | X |
| MAFFT(default) | 0.207 | 0.289 | 0.320 | 0.312 | 0.298 | OOM |
| PASTA | 0.184 | 0.242 | 0.344 | 0.323 | 0.245 | 0.206 |
| MAFFT-sparsecore | 0.112 | 0.230 | 0.303 | 0.281 | 0.267 | OOM |
| FAMSA | 0.212 | 0.262 | 0.232 | 0.220 | 0.253 | 0.185 |
| MAGUS | 0.087 | 0.236 | 0.276 | 0.253 | 0.171 | *0.182 |
| SALMA(all) | 0.090 | 0.232 | 0.312 | 0.291 | 0.168 | **0.175 |
| WITCH(all) | 0.102 | 0.234 | 0.339 | 0.312 | 0.170 | X |
| MAGUS+UPP(all) | 0.105 | 0.236 | 0.339 | 0.314 | 0.170 | **0.175 |
| | | average SPFP | | | | |
| Clustal-Omega | 0.285 | 0.221 | 0.234 | 0.233 | 0.251 | 0.359 |
| MUSCLE | 0.284 | 0.288 | 0.580 | X | X | X |
| MAFFT(default) | 0.182 | 0.247 | 0.214 | 0.210 | 0.155 | OOM |
| PASTA | 0.174 | 0.224 | 0.169 | 0.167 | 0.172 | 0.181 |
| MAFFT-sparsecore | 0.138 | 0.220 | 0.204 | 0.195 | 0.160 | OOM |
| FAMSA | 0.235 | 0.281 | 0.241 | 0.227 | 0.236 | 0.182 |
| MAGUS | 0.109 | 0.169 | 0.110 | 0.107 | 0.110 | *0.157 |
| SALMA(all) | 0.115 | 0.188 | 0.112 | 0.116 | 0.110 | **0.146 |
| WITCH(all) | 0.113 | 0.183 | 0.112 | 0.116 | 0.112 | X |
| MAGUS+UPP(all) | 0.111 | 0.181 | 0.110 | 0.113 | 0.112 | **0.146 |
| | | average runtime (hours) | | | | |
| Clustal-Omega | 0.352 | 0.016 | 0.223 | 0.199 | 0.286 | 0.663 |
| MUSCLE | 0.485 | 0.005 | 2.093 | X | X | X |
| MAFFT(default) | 0.013 | 0.001 | 0.064 | 0.136 | 0.592 | OOM |
| PASTA | 1.399 | 0.638 | 2.347 | 2.331 | 4.198 | 11.196 |
| MAFFT-sparsecore | 0.459 | 0.237 | 0.268 | 0.359 | 2.113 | OOM |
| FAMSA | 0.007 | 0.001 | 0.010 | 0.011 | 0.031 | 0.138 |
| MAGUS | 1.248 | 0.185 | 1.409 | 1.476 | 8.841 | *3.325 |
| SALMA(all) | 0.753 | 0.141 | 2.089 | 2.066 | 9.840 | **19.675 |
| WITCH(all) | 1.118 | 0.141 | 4.336 | 4.301 | 21.429 | X |
| MAGUS+UPP(all) | 1.060 | 0.139 | 2.780 | 2.882 | 10.013 | **19.959 |

**Table S4: Experiment 2: Average SPFN, SPFP, and runtime in hours for all tested methods on biological datasets. CRW refers to the five selected CRW datasets. 10AA is a set of ten biological protein datasets. Homfam(10) refers to the ten largest Homfam datasets, while Homfam(8) excludes the largest two, rvp and zf-CCHH, because MUSCLE failed to run on them. "OOM" denotes that the method encountered out-of-memory issues before hitting the runtime limit, and "X" denotes that the method failed to run or did not finish within the time limit. "(all)" for SALMA, WITCH, and MAGUS+UPP denotes that they use all full-length sequences to form the backbone alignment. *MAGUS was run in recursive mode on the Res dataset. **The MAGUS backbone for the Res dataset was run in recursive mode.**
